# Functional Characterization of an Electromagnetic Perceptive Protein

**DOI:** 10.1101/2020.10.07.329946

**Authors:** Sunayana Mitra, Carlo Barnaba, Jens Schmidt, Galit Pelled, Assaf A. Gilad

**Affiliations:** Department of Biomedical Engineering, Michigan State University, East Lansing MI, USA; Division of Synthetic Biology and Regenerative Medicine, Institute for Quantitative Health Science and Engineering, Michigan State University, East Lansing, MI, USA; Department of Obstetrics, Gynecology, and Reproductive Biology, Michigan State University, East Lansing, MI, USA; Division of Neuroengineering, Institute for Quantitative Health Science and Engineering, Michigan State University, East Lansing, MI, USA; Department of Radiology, Michigan State University, East Lansing, MI, USA

**Author notes:** Corresponding Author: **Assaf A. Gilad**.

**Keywords:** membrane protein, molecular imaging, magnetoreception, HaloTag, Total Internal Reflection Fluorescence (TIRF), *Kryptopterus vitreolus*

## Abstract

Magnetoreception, the response to geomagnetic fields is a well described phenomenon in nature. However, it is likely that convergent evolution led to different mechanisms in different organisms. One intriguing example is the unique Electromagnetic Perceptive Gene (EPG) from the glass catfish *Kryptopterus vitreolus*, that can remotely control cellular function, upon magnetic stimulation in *in-vitro* and *in-vivo*. Here, we report for the first time the cellular location and orientation of the EPG protein. We utilized a differential labelling technique to determine that the EPG protein is a membrane anchored protein with an N-terminal extracellular domain. The kinetics and diffusion dynamics of the EPG protein in response to magnetic stimulation was also elucidated using single particle imaging and tracking. Pulse chase labelling and Total Internal Reflection Fluorescence (TIRF) imaging revealed an increase in EPG kinetics post magnetic activation at a single particle level. Trajectory analysis show notably different EPG protein kinetics before and after magnetic stimulation in both 2 (free vs bound particle) and 3 state (free vs intermediate vs bound particle) tracking models. This data provides additional information to support and understand the underlying biophysical mechanisms behind EPG activation by magnetic fields and provides evidence for the basis of magnetoreception in the EPG protein that will aid in future studies that seek to further understand this novel mechanism. This study is important for understanding magnetoreception as well as developing new technologies for magnetogenetics – the utilization of electromagnetic fields to remotely control cellular function.

**Table of Contents Graphic:** 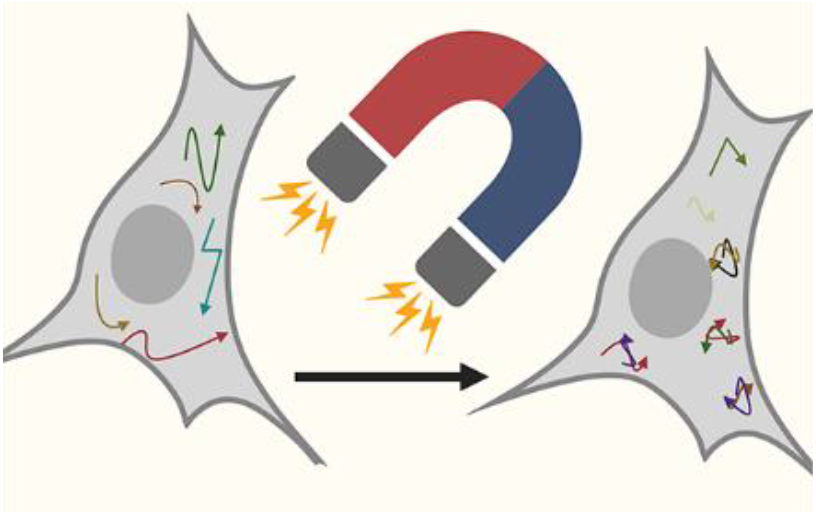

## Introduction

Ample evidence suggests that many organisms possess the ability of geomagnetic sensing. Nevertheless, for many animals the exact biophysical basis is elusive and there is limited information on this phenomenon(1,2). Though organisms like birds(3) and even recently humans(4) were explored, the majority of the research was done in fish, to explain migraton behavior and predator evasion(5), with recent examples in species such as zebrafish and medaka(6) as well as glass catfish(7) and eel(8). While many of the studies provide a holistic approach, very few looked into the molecular mechanism.

We have recently cloned a novel *Electromagnetic Perceptive Gene (EPG)*(9) from the glass catfish *Kryptopterus vitreolus*(10). When expressed in mammalian cells, the EPG shows a remarkable response to changes in electromagnetic fields (EMF) at the 50-150 mT range. This response is manifested by an increase in intracellular calcium(9,11). This gene is currently being explored in our lab as it has immense potential in the cellular modulation and could be incorporated into treatments of brain disorders(12). The importance of understanding this protein is growing in light of the interest in the potential of the expanding field electromagnetic and radiofrequency waves as a tool for remote cellular activation(13-16). Therefore, it is important to study the structure and function of proteins such as the EPG.

Here we sought to use imaging and particle tracking techniques to understand the functional properties of the EPG protein. Understanding the cellular localization and properties of this protein would give us greater insight into the nature and function of this gene. We opted to use single particle tracking method to follow our protein during various phases of activity, especially in tracking EPG activity pre and post magnetic stimulation. Single particle tracking techniques have historically used sensitive fluorescence microscopy techniques to allow particles of interest to be tracked during various stages of cellular uptake and transport(17). The advent of Total Internal Reflection Fluorescent (TIRF) microscopy has permitted very detailed visualizations of biological processes in real time(18).

We report for the first time new findings that determine the membranal orientation of the EPG protein. In addition, the successful use of TIRF based single particle tracking provides indication of clustering of th e EPG molecules upon magnetic stimulation. Thus, this study is another step in understanding some of the pieces of this puzzle, which eventually will allow better understanding of magnetoreception in fish as well as transformation of this unique ability into a useful scientific tool.

## Results & Discussion

### EPG is a membrane anchored extracellular protein

It has been previously shown that the expression of EPG was localized to the cellular membrane in HEK293 cells(11). The next meaningful question to answer is determining the orientation of EPG on the cellular membrane. Information about the localization and orientation is a critical step in further determining the kinetics and folding functions of the protein.

We continued the effort to determine the cellular localization of EPG. To that purpose, we constructed several vectors in which the EPG is fused to HaloTag®. HaloTag is a self-labeling protein tag, 297 residue peptide (33 kDa) that can be manipulated to bind to a synthetic ligand and fuse to a protein of interest(19). The advantage of using HaloTag fluorescent ligands is the ability to use a single genetic construct for multiple applications(20). To confirm the cellular orientation of the EPG protein across the membrane, we utilized a two-pronged approach with HaloTag and single particle tracking (SPT) technique. We used two different constructs with two different fluorescent probes, the cell permeable JaneliaFluor 646 (JF646) and the cell impermeable AlexaFluor 488 (AF488) that can bind the EPG-HaloTag fusion protein. HeLa cells were transfected with different EPG-HaloTag construct plasmid. The cells were then labelled using the AF488 and JF646 dyes 24 hours post transfection and imaged using fluorescent microscopy.

Figure **1A & B** show fluorescent images with the corresponding schematics below for HeLa cells transfected with EPG-HaloTag fusion construct with the tag at the N-terminus of EPG (Halo-N_term_-EPG; **Supplementary Figure S1**) and labelled with AF488 and JF646. Both the cell permeable (JF646 in yellow) and impermeable dye (AF488 in green) constructs showed robust fluorescence in HeLa cells. Alternatively, when we used EPG-HaloTag fusion constructs with the HaloTag at the C-terminus (EPG-C_term_-Halo; **Supplementary Figure S2**), only the cell permeable dye (JF646) showed successful labeling **(Figure 1D)**. There was no labeling in cells treated with the cell impermeable dye (AF488) (**Figure 1C)**. This shows that EPG protein is a transmembrane anchored protein that is oriented with its N-terminal end facing the extra cellular space and the C-terminal end being the membrane anchor, as illustrated by the schematics (**Figure 1E-H**). The strong fluorescent signal in from JF646 in both types of transfected cells also indicated robust expression of the respective EPG-HaloTag fusion proteins. These results also conform our previous finding that the EPG protein has a Ly6 motif that is found on extracellular region(9). This is an important key to unlocking further information about the downstream processes that EPG might be involved in.

**Fig. 1.**
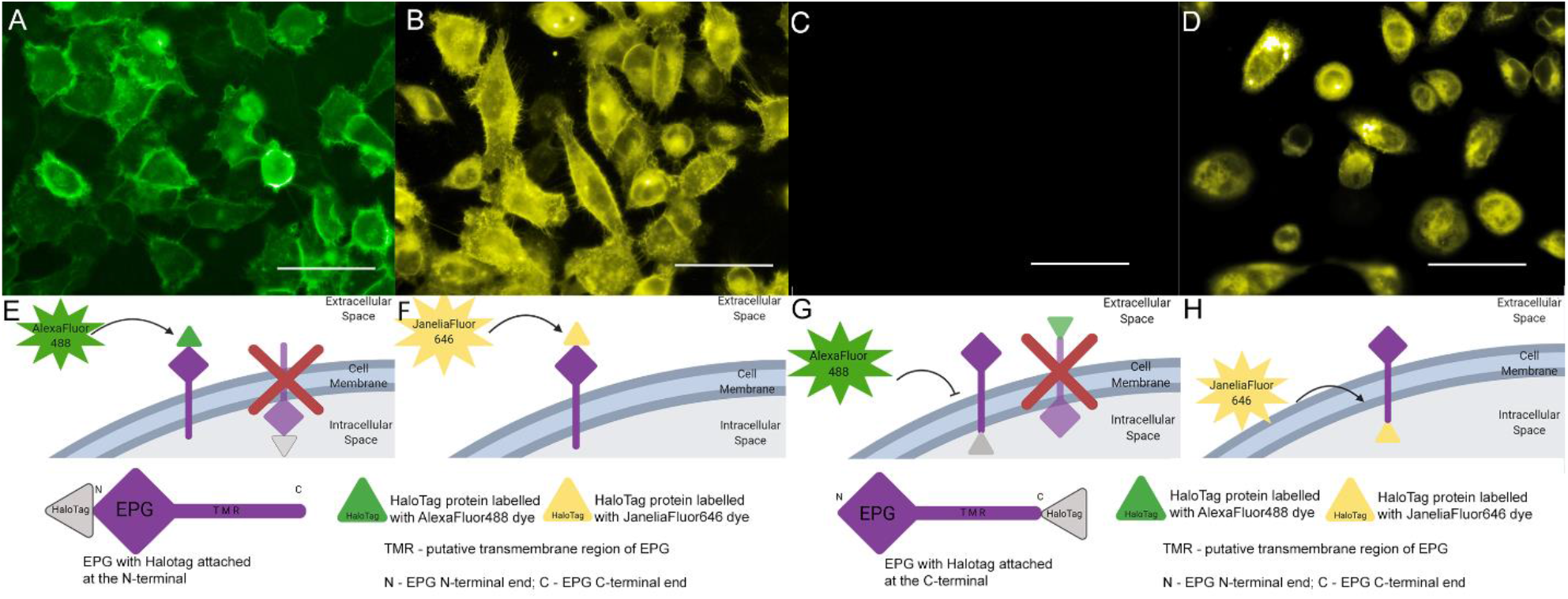
N-terminal and C-terminal HaloTag-EPG fusion protein dye labelling in HeLa cells. (**A** & **C**) images of the cells labelled with cell impermeable AlexaFluor488 dye (green) through the GFP filter. Only cells that express the HaloTag fused to N terminus were stained (**A**) but not cells that express the HaloTag fused to C terminus. These findings indicate that EPG is expressed on the membrane facing the extracellular space as depicted in the illustrations (E & G respectively). (**B** & **D**) images of the transfected cells labelled with cell permeable JaneliaFluor646 dye (yellow) and imaged through Cy5 filter. As illustrated in (**F** & **H**) this dye can label the protein in both orientations and used to demonstrate that both constructs, N terminus and C terminus fusion were expressed in sufficient level. Scale bar indicates 50µm.

### EPG molecular dynamics pre and post magnetic stimulation were probed using Pulse Chase labeling and TIRF microscopy

EPG protein has been established as being responsive to both static and electromagnetic stimulation in a variety of *in-vivo* and *in-vitro* models(9,11). To elucidate EPG’s mechanism, we sought to study the single particle dynamics of EPG’s response to magnetic fields using TIRF microscopy. TIRF microscopy solves the age old problem with out-of-focus fluorescence by restricting excitation parameters to a very small layer adjacent to the coverslip(9). This makes it possible to perform single particle detection at a high resolution. TIRF microscopy with live cell imaging could also be used to characterize the intracellular dynamics of our gene of interest(9).

HeLa cells were transfected with the construct expressing Halo-N _term_-EPG. Pulse chase labelling was performed 24 hours post transfection, by initially flooding the transfected cells with the cell impermeable dye (AF488). This cell impermeable ligand labels the entire cell surface pool of Halo-N_term_-EPG molecule **(Figure 2A)**. We subsequently labeled these cells with the secondary, cell permeable JF646 dye. The JF646 cell permeable dye selectively labels only the intracellular reserve of Halo-N_term_-EPG molecules **(Figure 2D)**. This was done to ensure a reasonable number of fluorescent particles could be detected at the surface of the cell during the subsequent TIRF microscopy and analysed by particle analysis sioftware without overwhelming it. That is an important consideration to ensure that a particle tracking software used for image analysis post acquisition is not overwhelmed. Next, the labelled cells were subjected to a static magnetic field of 50 mT for 10 seconds **(Figure 2B and E)**, the magnet was removed and the cells were allowed to rest for 10 mins before being imaged again (**Figure 2C and F**). **Supplementary Video SV1 (A-F)** show single molecule imaging dynamics of cells pulse labelled with AF488 and JF646 by high resolution TIRF microscopy before, with and after 10 mins of magnetic stimulation. The images in the top panel show strong overall fluorescence under all conditions indicating no discernable changes within this pool of AF488 labelled molecules. However, in the bottom panel of JF646 labelled cells we can see individual EPG-HaloTag fusion protein molecules as bright spots easily distinguishable from the background. These results show that TIRF microscope can be used successfully to observe single particle dynamics of EPG-HaloTag fusion, at single particle level, before and after magnetic stimulation. This is consistent with TIRF assays that have been successfully used to quantify fluorescently tagged proteins at concentrations in the pico to nano molar range(21). These results also show that magnetic stimulation likely leads to recruitment of EPG molecules to the surface of the cell.

**Figure 2.**
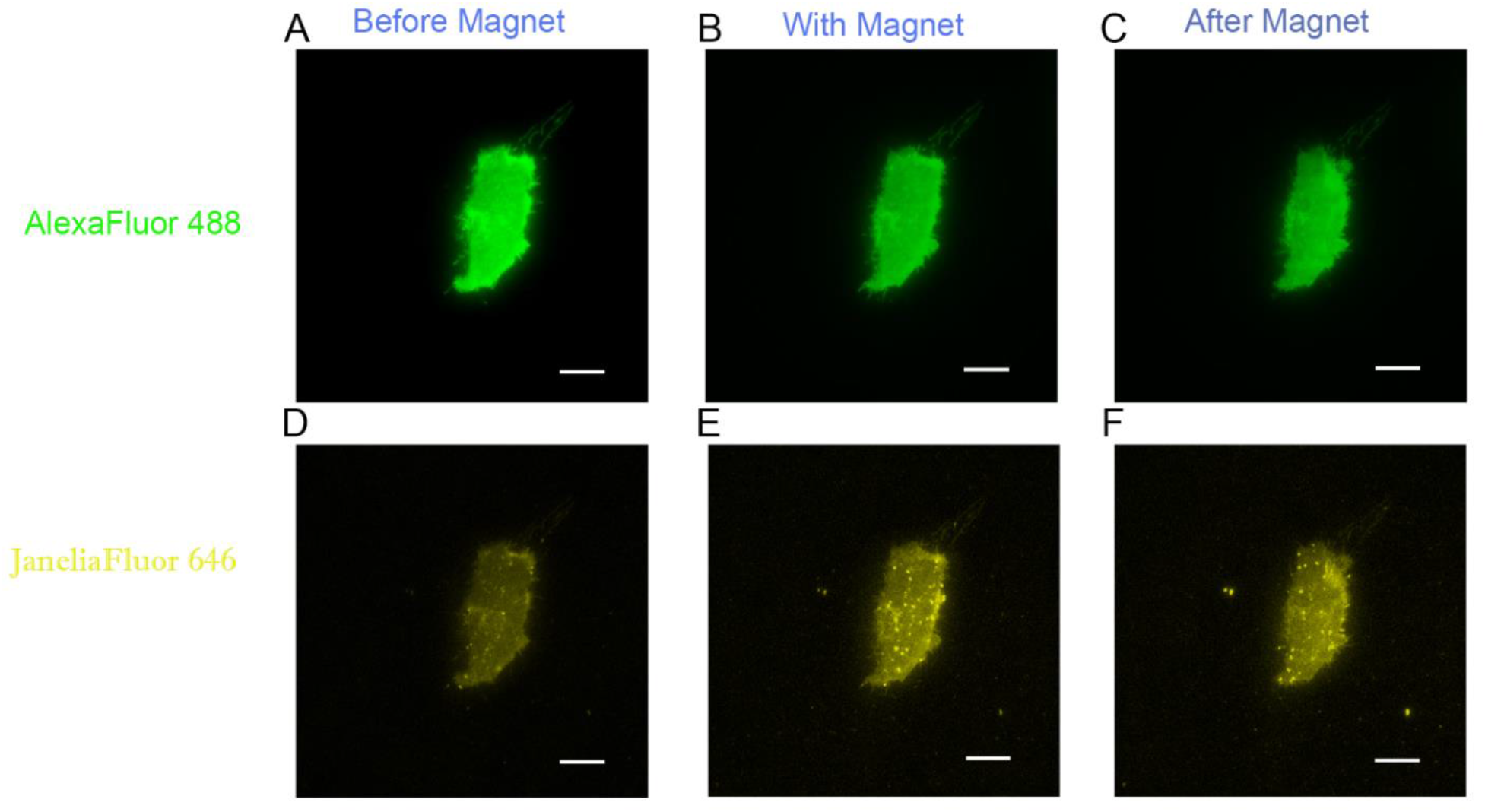
Single molecule imaging before and after magnetic stimulation in labelled EPG. (**A**) shows HeLa cells transfected with EPG pulse labelled with AF488 (green) followed by JF646 (red), before magnetic stimulation. The labelling is shown to be more global with AF488 due to high expression of proteins on the surface by transiently transfected cells. (**B**) shows single molecule imaging in both AF488 and JF646 dye labelled cells with images during magnetic stimulation using a static magnet (50 mT). (**C**) Shows images of the same cells with the same dyes but 10 mins after removal of magnetic stimulation. No change is seen on the AF488 labelled surface pool over different conditions. Gradual increase in cell surface appearance of intracellularly JF646 labeled particles is seen. Scale bar 10 µm.

### Single particle tracking and localization detects real time trajectories of EPG molecules

To track EPG particle dynamics in real time before and after magnetic stimulation, we implemented a MATLAB based single particle tracking (SPT) script that was developed previously and used to generate dynamic maps of particles that were tracked with high resolution for epidermal growth factor(22). We used this technique to generate individual location and trajectories of EPG particles alternating between free and confined states. **Figure 3** shows an image of the same single cell taken at 2 seconds into the recording either before magnetic induction (**A**) or with magnetic stimulation (**B**) or ten minutes after termination of magnetic stimulation (**C**). **Supplementary Video SV2 (A-C)** show the results of frame by frame tracking of EPG particle trajectories before, with and after 10 mins of magnetic stimulation.

**Figure 3.**
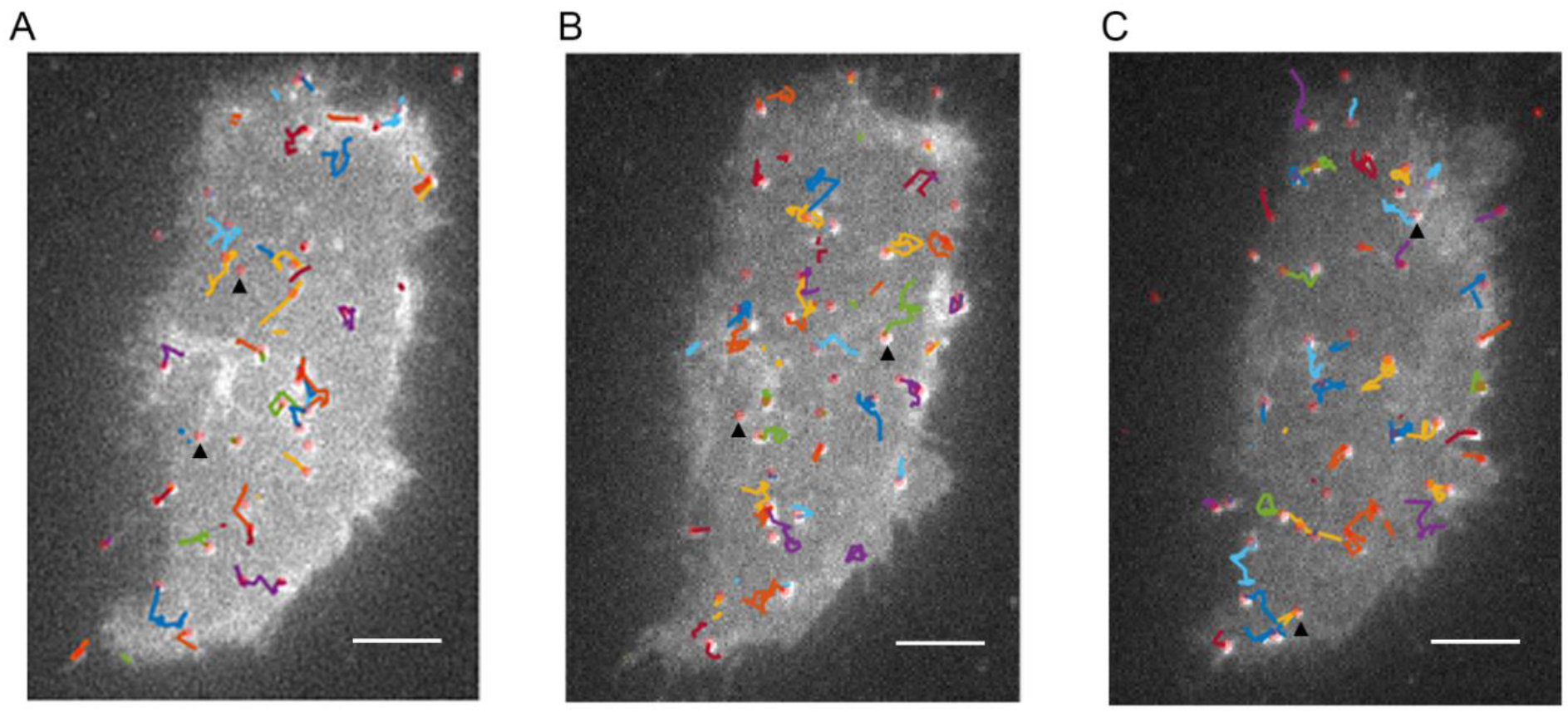
Tracking single EPG molecules in live cells. A representative image of a single HeLa cell expressing HaloTag-N _term_-EPG labeled with JaneliaFluor 646 labeled. (A) before (B) during and (C) after magnetic stimulation. Black arrowheads represent individual tracked Halo-N_term_-EPG molecules. Individual particle trajectories are represented by different colored tracks. Scale bar 5 µm.

### Trajectory analysis algorithm shows notably different EPG protein kinetics before and after magnetic stimulation in both 2 and 3 state models

We used particle tracking of EPG trajectory and generated data of pooled trajectories before, during and after magnetic stimulation. We sought to use a fit model to assess displacement histograms of the EPG trajectories and generate diffusion coefficients of particles. We used the generated SPT data and analyzed it using Spot-On, an algorithm(23) developed to track particle data based on bound vs unbound or free state configurations. Spot-On uses SPT information and generates data to infer the size of a subpopulation of molecules. Spot-On assumes that the target protein can exist in different sub-populations, with different diffusion coefficients either in two or three different states(24). Furthermore, this enables us to establish changes in particle behavior (based on diffusion coefficients) before and after magnetic stimulation. **Figure 4** shows data comparison between bound vs free (two-state) diffusion kinetics of EPG particles before, during and 10 mins after magnetic stimulation. The two-state model assumes that the particles tracked exist in two distinct populations, either free or bound. In this case the bound EPG particles could be the ones forming multimers and/or interacting with other proteins.

**Figure 4:**
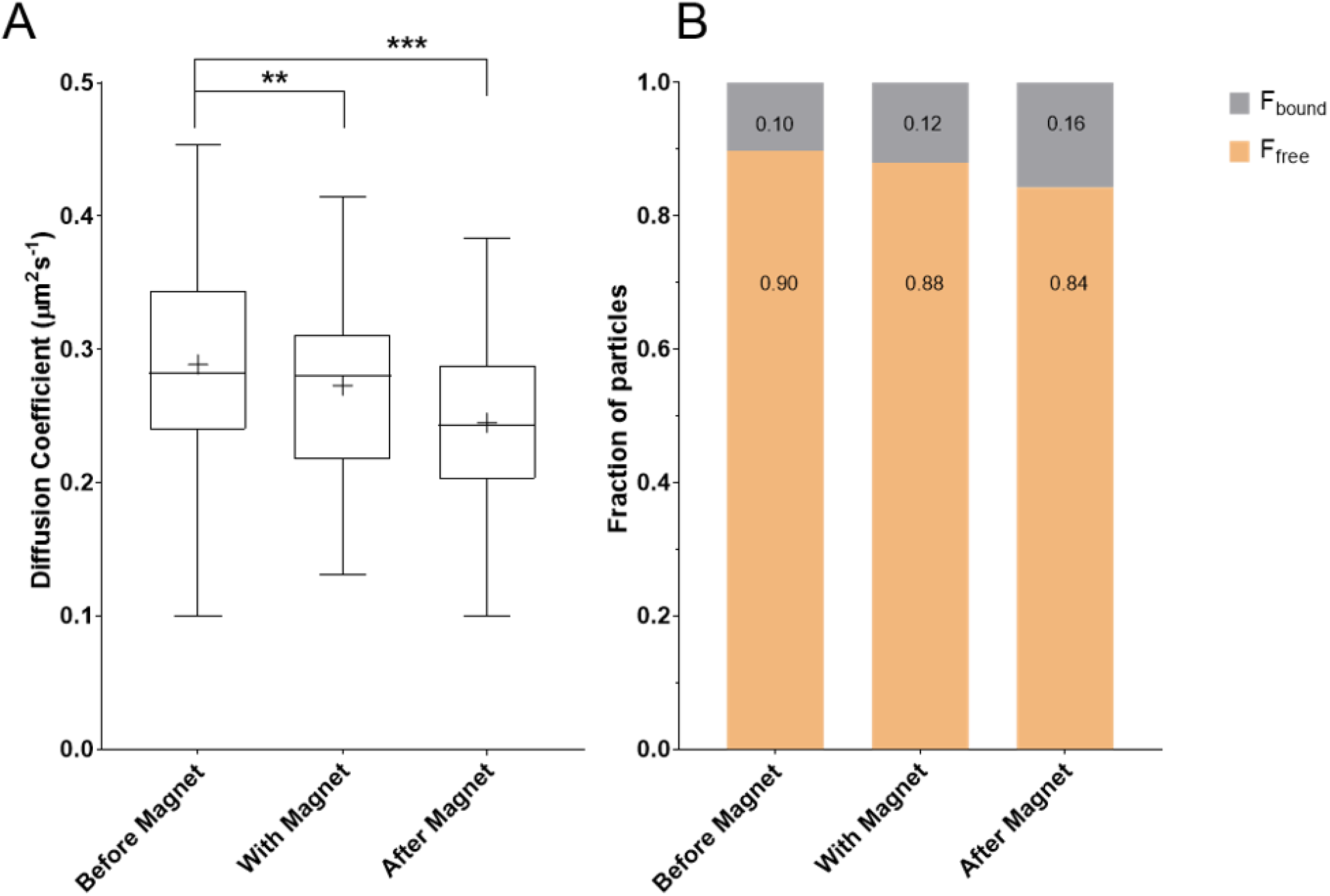
Two-State model shows different diffusion kinetics between pre and post magnetic stimulation. **(A**) A box and whisker plot shows comparison of combined particle diffusion coefficients of free particles (D_free_) and **(B)** fraction of particle population in either free, F_free_ or bound, F_bound,_ under different conditions. There is a significant decrease in D_free_ upon magnetic stimulation (* = p < 0.05, *** = p < 0.0001) with a corresponding decrease population of free particles F_free_. Conversely magnetic stimulation causes increase in bound particles. Whiskers are from minimum value to the maximum, the box extends from 25 ^th^ percentile to the 75^th^ percentile and the central line in the box is plotted at the median value. Means are marked by ‘+’. n= 5 independent experiments with 4-9 individual cells in each experiment

Figure **4A** **and B** show free state diffusion coefficients and the fraction of particles in the free and bound states under different conditions. The effect of magnetic stimulation on EPG particle diffusion coefficients and the population fractions were analyzed using two-tailed, paired t-test. The Before Magnet group was compared to the With Magnet group as well as to the After Magnet group in all cases. There is a gradual decrease in the diffusion coefficients of the free particles, D_free,_ with a corresponding decrease in the population of free particles, F_free_. The D_free_ value was found to be significantly decreased from what was seen Before Magnet as compared to both With Magnet (p= 0.0075) and After Magnet (p<0.0001) (**Figure 4 A**). This suggests that there aree increased interactions of EPG molecules with other proteins or formation of multimers upon magnetic stimulation. These interactions also result in slowing of the protein molecules, with more and more molecules transitioning to the bound state from the free state. This is also shown by an increase in the bound population, F_bound_, while under magnetic stimulation and after, as compared to unstimulated particles (**Figure 4 B**). There is an overall slowing of the bound particles as well (**Supplementary Figure S4** (p= n.s)), but the effect is very small as the actual diffusion coefficients of the bound population, D_bound_ is very small to begin with. We believe this is because the bound EPG particles are very slow due to protein-protein interactions.

Based on the previously analyzed two state model data we sought to extend this to a three state model that assumes that EPG exists in three states, a Free and Fast (F_free1_) state, a Bound state (F_bound_) and an intermediate, Free and Slower, (F_free2_) state (**Figure 5A and B**). Here too, there is a gradual decrease in the diffusion coefficients of free population, showing a significant difference in D_free_ of the After Magnet population as compared to D_free_ of Before Magnet population (p< 0.0001) (**Figure 5A**), with the corresponding decrease in population of F_free1_ particles similar to what is seen in the F _free_ population as in the two-state model. Under the three-state assumption, the F_bound_ EPG particles are close to truly immobile where they might have assumed the final configuration (with self or other proteins) under magnetic stimulation and thus do not show much difference in diffusion coefficients under different conditions. This is indicated by the unchanging D_bound_, even though there is an increase in the F_bound_ molecules (**Figure 5 B** and **Supplementary Figure S5 A**). The intermediate F_free2_ molecules represent the population of EPG molecules in the process of getting associated with each other or other proteins due to the magnetic stimulation. The F_free2_ molecules shows an increase with stimulation, perhaps due to more molecules transitioning form Free1 to the Free2 state, with a continuing decrease in the correponding D_free2_, as the dynamics decrease (**Figure 5 B** and **Supplementary Figure S5 B**). These results show that magnetic stimulation may lead to changes in the way EPG molecules interact with the dynamic environment. The possibility of EPG particle forming multimers or associating with other proteins gives new insights into EPG mechanism.

**Figure 5:**
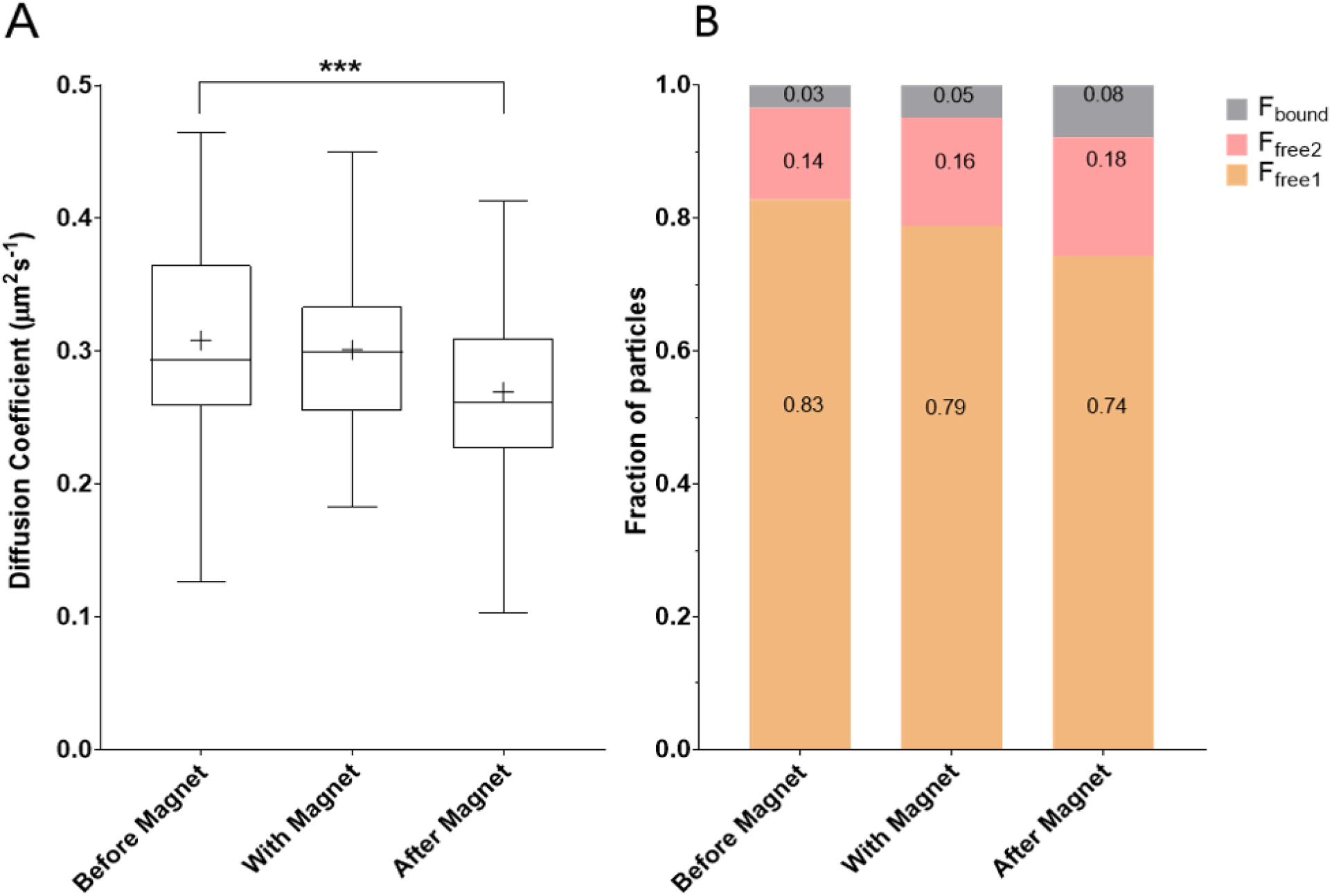
Three-State model show different diffusion kinetics between pre and post magnetic stimulation. This figure shows comparison of free particle diffusion coefficients D _free1_(**A**) and fraction of particle population (**B**) in either free, F_free1_, intermediate, F_free2_ or bound, F_bound_, under different conditions. There is a significant decrease in D _free1_ upon magnetic stimulation (*** = p < 0.001) with a corresponding decrease in population of free particles, F_free1_. Magnetic stimulation causes increase in both intermediary F _free2_ and bound F_bound_ particles. Whiskers are from minimum value to the maximum, the box extends from 25 ^th^ percentile to the 75^th^ percentile and the central line in the box is plotted at the median value. Means are marked by ‘+’. n= 5 independent experiments with 4-9 individual cells in each experiment.

We used spot-on to implement 2 and 3 state models to track EPG particle trajectories as a measure of diffusion coefficients. It is simplistic to state that the three-state model is better due to its containing two additional analysis parameters, we found that the two-state model has shown to be sufficient as well. Historically the two-state model has been used for transcription factors and simple protein interactions while the three-state model has been shown to be ideal for including particles that are in active state of diffusion and cellular confinement. Recently SPT has been used to track membrane proteins bound to fluorescent magnetic nanoparticles (FMNPs) and manipulated with a magnetic needle(25). Here we have shown tracking of unique EPG particles with inherent magneto-sensing ability, under magnetic stimulation with a static magnet.

The race for novel modulation techniques that can non-invasively, remotely control cellular function led to breakthrough methods. One such technique that gained considerable ground has been optogenetics that used light to control cellular function(26-28). Our lab pioneered another novel technique that uses magnetic stimulation to activate EPG protein to control cellular function(9). In clinically relevant rodent model of peripheral nerve injury, the EPG demonstrated to evoke neural responses when stimulated with electromagnetic field and consequently can restore cortical excitability(12). Moreover, the EPG can be stimulated with magnetic nanoparticles as well(11). These studies indicate the need of developing the EPG technology even further, and to that end, a deep mechanistic and structural understanding of the EPG was required.

The EPG was cloned from the fish, *Kryptopterrus vitreolus*, and show very little homology for any known gene. Recently, another novel gene from zebrafish was cloned. The gene termed “Bouncer” showed the closest homology to the EPG and was also found to be Ly6/uPAR protein and is a crucial fertilization factor in the zebrafish(29). Nevertheless, very little is known about the EPG function in its natural environment or when expressed in mammalian cells. Therefore, in this study we started by elucidation of the membrane orientation of the EPG. We report for the first time the subcellular localization of the EPG protein which is embedded in the plasma membrane facing the extracellular space. The localization of EPG on the membrane implies that protein-protein interactions, important to the EPG’s function occur on the outer side of the membrane.

By designing a “pulse chase” experiment using dual dyes with TIRF microscopy we were able to show changes in the kinetics of the EPG protein following magnetic stimulation. There is evidence pointing to EMF stimulation resulting in increasing the number of the EPG molecules on the surface of the membrane. In addition, we provide a mechanistic model showing different kinetics, using single particle tracking, which indicates increased interactions of EPG molecules with itself or other proteins or formation of multimers upon magnetic stimulation.

In conclusion, we have reported success in the first step of deciphering the function of EPG at the cellular level. Deciphering the structure and cellular localization of this gene/protein will provide useful insights in elucidating the downstream processes that are involved in cellular control. Additional research is required in the future to assess EPG folding mechanisms and other post translational mechanisms, as well as transmembrane and intracellular signaling pathways. Nevertheless, the findings described in this study are important by providing guidelines for evolving the next generation of EPG based technologies of genetically encoded cellular modulators.

## Experimental Procedures

### HeLa cell culture

HeLa cells were grown with DMEM media (Invitrogen) supplemented with 10 % FBS and 1% PenStrep solution (Invitrogen) and maintained in 25 cm^2^ polystyrene flasks. For experiments, HeLa cells we plated in 24 well plates (Cellvis) at a density of ∼ 0.5 × 10^5^ cells per well and grown for 24 hours prior to transfection.

### EPG Cloning and Transfection

The EPG open reading frame (ORF), as described in(9) contains a putative 20 amino acid membrane signal sequence at the N terminal region predicted using SignalP 5.0, a neural networks based online signal peptide prediction program(30) (**Supplementary Fig S3**). We designed sequence specific primers to generate separate DNA sections of the signal sequence and rest of functional EPG fragment using PCR. The NEBuilder HiFi DNA Assembly kit (New England Biolabs) was used to combine the two split EPG sections with N-Terminal HaloTag Plasmid PHTN (Promega) such that the N-terminal HaloTag sequence is preceded by the signal sequence at its 5’ end and succeeded by the rest of the EPG sequence at the 3’ end, creating the Halo-N_term_-EPG construct. To create the EPG-C_term_-Halo construct, the EPG fragment was cloned upstream to the HaloTag region on the C-Terminal Tagged HaloTag Plasmid, PHTC (Promega) using the same kit as above.

HeLa cells were transiently transfected with either the Halo-N_term_-EPG or the EPG-C_term_-Halo constructs using the Lipofectamine 3000 transfection kit (Invitrogen) following the kit instructions. Transiently transfected cells have very high levels of protein expression due to the introduction of several copies of the DNA vector into the cells. To control the level of protein expression so that subsequent HaloTag labelling results in a finite number of individually labeled protein molecules we used only 10 ng of DNA /well for transfection reactions.

### HaloTag Single Dye Labelling

HeLa cells were labeled with appropriate HaloTag reagents 24-hours post transfection as per manufacturer instructions. Briefly, for labelling with AF488, 1µM of the dye conjugated HaloTag ligand was added to each well of 24-well plate of transfected cells. Cells were then incubated for 15 minutes at 37°C and 5% CO_2_ in a cell culture incubator. The cells were then rinsed twice with pre-warmed complete media, to wash out excess ligand. Finally, prewarmed Fluorbrite media (Invitrogen) supplemented with 10% FBS was added in the wells for fluorescent microscopy. For labelling with JF646, 200nM dye conjugated ligand was used per well. After similar incubation as above, the cell media was simply replaced with prewarmed, supplemented Fluorbrite and wells imaged.

### HaloTag ‘Pulse-Chase’ Labelling

HeLa cells, transfected with Halo-N_term_-EPG construct, were labeled 24-hours post transfection. The ‘pulse’ label, AF488 conjugated ligand, was added at a concentration of 500nM per well. The cells were then incubated as described above. Following the pulse incubation, the ‘chase’ label, JF646, was added to the same wells at a concentration of 100 pM per well. Such low concentration of the chase label is necessary to ensure just enough protein molecules are labelled as to not to overwhelm the subsequent particle analysis programs. The cells were incubated for 30 seconds and then washed twice with prewarmed media, ending with prewarmed, supplemented Fluorbrite.

### Live Cell Epifluorescent Microscopy

Labelled HeLa cells were initially imaged using the BZ-X770 automated microscope (Keyence). The 24-well plates were maintained at 37°C with 5% CO_2_ atmosphere using a dedicated stage top incubator STRF-KIW (Tokai-Hit). AF488 labelling was visualized by imaging through the GFP filter OP-87763 (Keyence) and a 20X Objective. JF646 labeled cells were visualized using the Cy-5 filter OP-87766 and the same objective.

### Live Cell Single Molecule Imaging

Labelled cells were imaged using an Olympus TIRF enabled microscope with four laser sources at 405nm, 488nm, 562 nm and 647nm, dual iXon Ultra 897 EMCCD cameras (Andor), a 100× oil-immersion objective (Nikon), and an environmental chamber to maintain temperature, humidity and CO_2_ levels. Cells of interest were initially imaged (before magnet dataset) simultaneously at 488 nm with 15% laser power, for AF488, and at 646nm with 20% laser power, for J646. Imaging was done continuously for 10 seconds with an exposure rate of 200 ms, to generate a rate of 7 frames per second. A static rare earth neodymium magnet was placed on the top of the cell plate cover to stimulate the cells with a 50 mT magnetic field. The cells were then imaged again as above without changing any setting to generate the With Magnet dataset. For the After Magnet dataset, the magnet was removed, and the cells were left undisturbed for 10 minutes and imaged again using the same setting as above. All imaging was carried out under HILO conditions(31) Image sets were exported onto ImageJ software for subsequent image visualization and analysis.

### EPG-Halotag Single Particle Tracking

EPG-Halotag TIRF image sets were analyzed with MATLAB R2020a (MathWorks Inc.), using the SLIMfast to generate single particle trajectories by utilizing the Multiple-Target Tracing (MTT) algorithm(22,32). To analyze single EPG particle diffusion, the maximal expected Diffusion Coefficient was set to 2 µm^2^s^-1^. The trajectories were then analyzed using Spot-On(23) for on trajectories two frames or longer.

The bound and free fractions and the corresponding diffusion coefficients were computed from the PDF curve over eight time intervals, considering a maxima of four jumps. The following settings were used on the Spot-On algorithm with the 2-state model: bin width 0.01, Timepoints 8, jumps to consider 4, use entire trajectories-No, Max jump (µm) 5.05. For model fitting the following parameters were chosen: D_bound_ (µm^2^ s^-1^) min 0.0001 max 0.1, D_free_ (µm^2^ s^-1^) min 0.15 max 5, Localization error (µm) fitted from data (min 0.01 max 0.075), and dZ (µm) 0.7, Model Fit PDF, Iterations 2, UseWeights-No. For the 3-states model, the only parameters changed were the diffusion limits, being D_bound_ (µm^2^ s^-1^) min 0.0001 max 0.01, D_free,1_ (µm^2^ s^-1^) min 0.1 max 5, D_free,2_ (µm^2^ s^-1^) min 0.01 max 0.5.

## Author Contribution

Conceptualization (S.M., J.S. G.P and A.A.G) Methodology (S.M., C.B., J.S. and A.A.G) Investigation (S.M.); Formal Analysis (S.M. and C.B) Writing – Original Draft (S.M. G.P. and A.A.G) Supervision (A.G.) Funding Acquisition (G.P. and A.A.G).

## Acknowledgements

We thank Rita Martin for manuscript editing.

## Funding Information

This project is supported in part by the NIH grants R01 NS098231 (AAG and GP), R01 NS104306 (AAG) and P41 EB024495 (AAG).

### Disclaimer

The content is solely the responsibility of the author s and does not necessarily represent the official views of the National Institutes of Health.

## Supporting Information

Supplementary Data: Includes figures of plasmid maps, EPG putative signal sequence and diffusion coefficient data D _bound_ and D_free2_. (from two as well as three state analysis).

Supplementary Movies: All movies were captured at and are played at 5fps

**Movie SV1 (A-C)**. Corresponding movie of **Fig. 2 A-C** Halo-N_term_-EPG particles are labelled with AlexaFluor 488 dye.

**Movie SV1 (D-F)**. Corresponding movie of **Fig. 2 D-F** Halo-N_term_-EPG particles are labelled with JaneliaFluor 646 dye.

**Movie SV2 (A-C)**. Corresponding movie of **Fig. 3 A-C** Halo-N_term_-EPG particles are labelled with JaneliaFluor 646 dye.

