## Supporting Information for "Functional Characterization of an Electromagnetic Perceptive Protein"

### Supporting Information
for

### Functional Characterization of an Electromagnetic Perceptive Protein

Sunayana Mitra[1,2], Carlo Barnaba[2,3], Jens Schmidt[2,3], Galit Pelled[1,4,5] & Assaf A. Gilad[1,2,5]

[1] Department of Biomedical Engineering, Michigan State University, East Lansing MI, USA

[2] Division of Synthetic Biology and Regenerative Medicine, Institute for Quantitative Health Science and Engineering, Michigan State University, East Lansing, MI, USA

[3] Department of Obstetrics, Gynecology, and Reproductive Biology, Michigan State University, East Lansing, MI, USA

[4] Division of Neuroengineering, Institute for Quantitative Health Science and Engineering, Michigan State University, East Lansing, MI, USA

[5] Department of Radiology, Michigan State University, East Lansing, MI, USA

Corresponding author:

Assaf A. Gilad Ph.D.


### Supplementary Figures

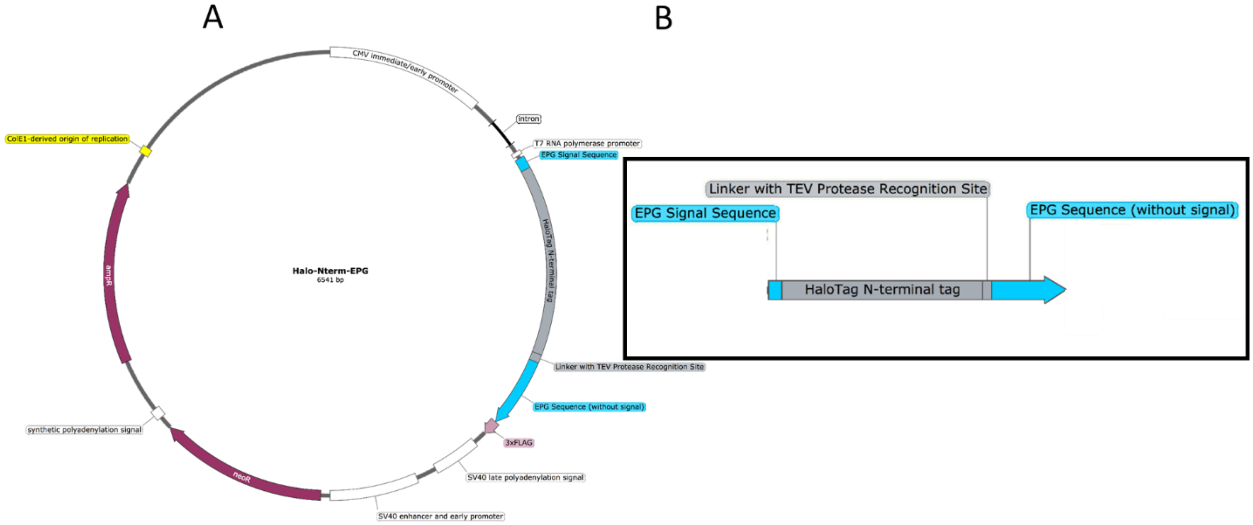


**Figure S1: Plasmid Map of Halo-N$_{term}$-EPG.** This figure shows entire plasmid map of the Halo-N$_{term}$-EPG construct (**A**) and the schematic of the EPG-HaloTag fusion gene (**B**). The EPG signal Sequence and the EPG protein sequences were generated from the same EPG gene using PCR. The entire construct was created using 2-step HiFi Assembly cloning kit (NEB). EPG protein sequence was inserted 3′ to the HaloTag in the first step and the EPG signal sequence was inserted in the second step 5′to the HaloTag. Plasmid map was created using SnapGene software (GSL Biotech).

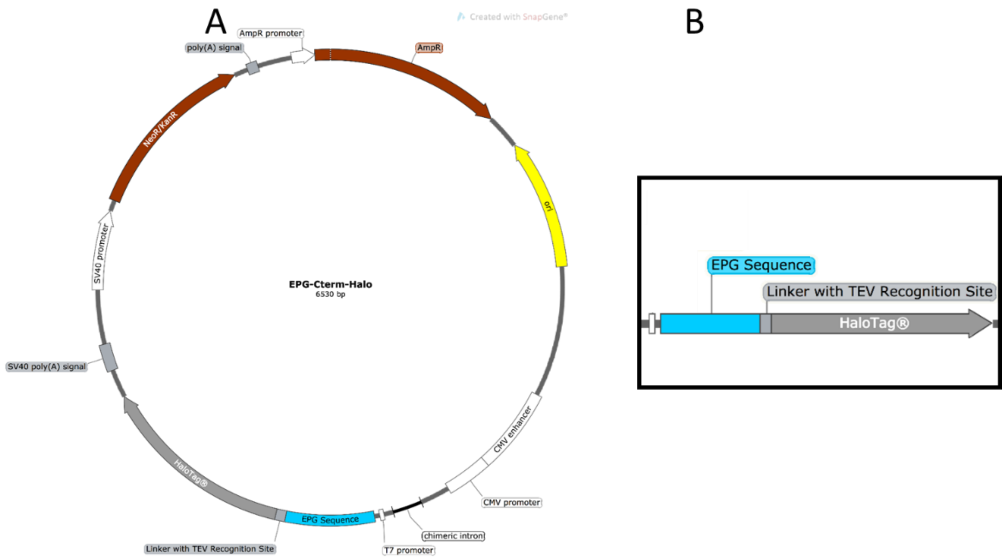


**Figure S2: Plasmid Map of EPG-C$_{term}$-Halo.** This figure shows entire plasmid map of the **EPG-C$_{term}$-Halo** construct (**A**) and the schematic of the EPG-HaloTag fusion gene (**B**). The EPG Sequence was generated from in house EPG plasmid using PCR. The construct was created using HiFi Assembly cloning kit (NEB). EPG sequence was inserted 5′ to the HaloTag. Plasmid map was created using SnapGene software (GSL Biotech).

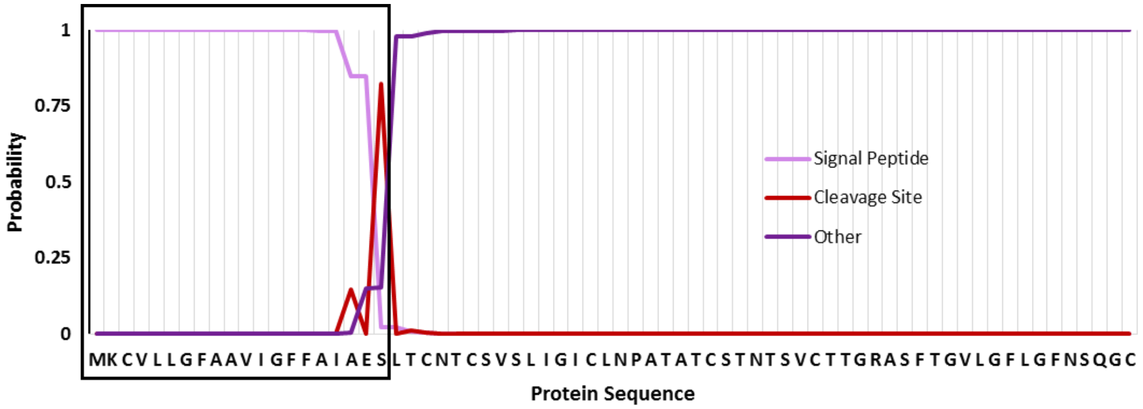


**Figure S3: Putative signal sequence of EPG.** This figure shows EPG amino acid sequence with the possible membrane signal sequence highlighted (black box). The program Signal P was used to predict the membrane signal peptide (mauve), the cleavage site (red) and protein sequence (purple).

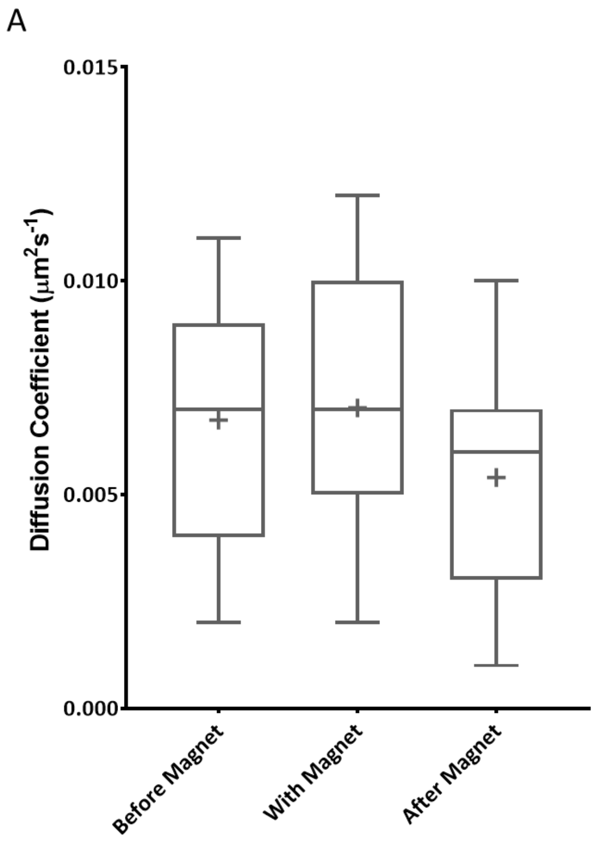


**Figure S4: Diffusion Coefficient of Bound EPG Particle Population using Two-State SPT analysis.** This figure shows the comparison of diffusion coefficient, $D_{bound}$, of the bound particle population under Before, With and After magnetic stimulation. An overall decrease is seen in $D_{bound}$ 10 minutes after removal of the magnet as compared to the diffusion coefficient before the introduction of the magnet.

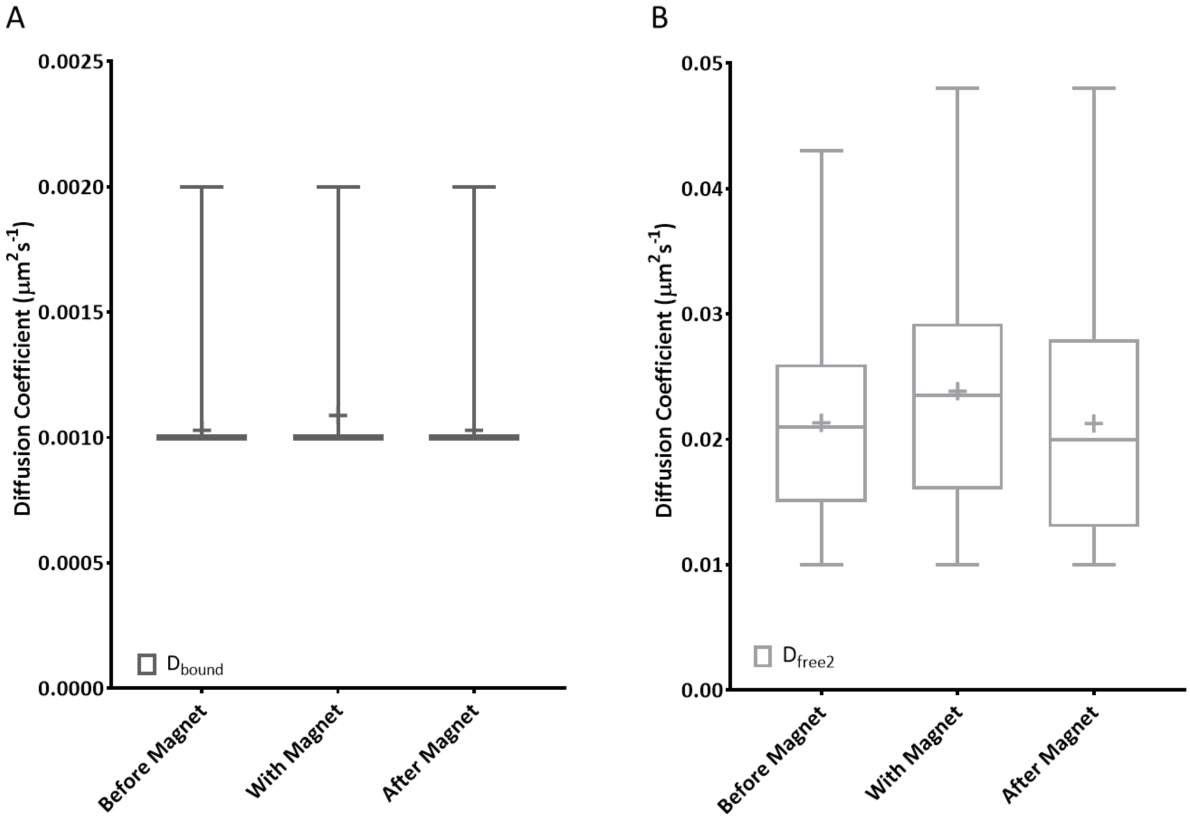


**Figure S5: Diffusion Coefficients of Intermediate and Bound EPG Particle Populations using three-State SPT analysis.** This figure shows the comparison of diffusion coefficient, **(A)** $D_{bound}$, of the bound particle population and **(B)** $D_{free2}$ of the intermediate particle population under Before, With and After magnetic stimulation. $D_{bound}$ is seen to be unchanged in the course of the experiment. An overall decrease is seen in $D_{free2}$ 10 minutes after removal of the magnet as compared to the diffusion coefficient before the introduction of the magnet.

### Supplementary Movies

All movies were captured at and are played at 5fps

**Movie SV1 (A-C).** Corresponding movie of **Fig. 2 A-C** Halo-$N_{term}$-EPG particles are labelled with AlexaFluor 488 dye.

**Movie SV1 (D-F).** Corresponding movie of **Fig. 2 D-F** Halo-$N_{term}$-EPG particles are labelled with JaneliaFluor 646 dye.

**Movie SV2 (A-C).** Corresponding movie of **Fig. 3 A-C** Halo-$N_{term}$-EPG particles are labelled with JaneliaFluor 646 dye**.**
